# TCR clustering by contrastive learning on antigen specificity

**DOI:** 10.1101/2024.04.04.587695

**Authors:** Margarita Pertseva, Oceane Follonier, Daniele Scarcella, Sai T. Reddy

**Affiliations:** Department of Biosystems Science and Engineering, ETH Zurich, 4058 Basel, Switzerland; Life Science Zurich Graduate School, ETH Zurich and University of Zurich, 8006 Zurich, Switzerland

**Keywords:** T cell receptor, contrastive learning, ESM model, deep learning, TCR clustering

## Abstract

Effective clustering of T-cell receptor (TCR) sequences could be used to predict their antigen-specificities. TCRs with highly dissimilar sequences can bind to the same antigen, thus making their clustering into a common antigen group a central challenge. Here, we develop TouCAN, a method that relies on contrastive learning and pre-trained protein language models to perform TCR sequence clustering and antigen-specificity predictions. Following training, TouCAN demonstrates the ability to cluster highly dissimilar TCRs into common antigen groups. Additionally, TouCAN demonstrates TCR clustering performance and antigen-specificity predictions comparable to other leading methods in the field.

## Introduction

T-cells play a pivotal role within the adaptive immune system, orchestrating molecular and cellular responses against pathogens and diseased cells (e.g., tumor cells) [1]. Each T-cell is characterized by a unique T-cell receptor (TCR) displayed on its surface, enabling the molecular recognition of pathogenic or diseased antigens in the form of intracellularly processed peptides loaded on to major histocompatibility complex (MHC) molecules [2]. TCRs consist of two chains: alpha and beta, which include variable regions (Vα, Vβ) possessing three hypervariable loops referred to as complementarity-determining regions (CDRα-or CDRβ-1, -2, -3) [3,4]. The CDRα and CDRβ loops come in contact with peptide-MHC complex, determining TCR specifcity. CDR3 loops are particularly diverse and have a major impact on peptide-MHC (pMHC) recognition [3,4]. TCR chain diversity is achieved through V(D)J recombination, which joins a random set of V, (D) and J gene segments together. During recombination, non-templated nucleotide changes are introduced into CDR regions [5], resulting in a highly diverse TCR repertoire, which has been theoretically estimated to be > 10^20^ [6]. It is estimated that a single pMHC can be recognised by hundreds to thousands of distinct TCRs [5], with some of these TCR Vα and Vβ sequences sharing common amino acid patterns (or motifs) in the CDRs[6,7]. However, it has also been shown that unique TCRs with specificity to the same pMHC antigen can also have highly dissimilar Vα / Vβ sequences [8]. Furthermore, TCRs are known to be cross-reactive by recognising multiple pMHCs. This trait is believed to be essential for covering all possible antigen diversity [9].

Clustering TCRs with the same antigen-specificity together is a major challenge in immunoinformatics. This is exacerbated by limited availability of public data, where there is only a fraction of human TCR repertoire and MHC allele diversity accessible (e.g., VDJdb or IEDB) [10]. These datasets are also highly unbalanced, with the majority of identified TCRs exhibiting specificity towards only 3-6 HLA alleles. For instance, approximately 20.7% of MHC I-restricted human TCRs target the HLA-A*02:01 allele in the VDJdb database. Furthermore, there is a notable imbalance in peptide antigen representation, with 97% originating from viral sources, and approximately 100 peptides accounting for 70% of all TCR-pMHC pairs [10].

Most TCR clustering and analysis tools have relied on Vβ-chain sequence information only, which is a consequence of the scarcity of paired-chain TCR (Vα and Vβ) data [10]. The recent emergence of single-cell sequencing is leading to the high-throughput generation of paired-chain TCR data [11,12], which are enabling improvements in TCR clustering analysis [13,14]. Current approaches to TCR clustering and pMHC binding specificity prediction can be broadly categorized into distance-based and feature-based methods [14]. Distance-based methods, such as TCRdist and GLIPH, explicitly define similarity metrics between two TCRs by quantifying factors such as number of amino acid differences between the CDR regions, position of the mismatched amino acids, and differences in CDR length [6,7]. In this context, the TCR similarity is determined without considering the antigen. Feature-based approaches employ machine learning algorithms to learn the underlying rules of TCR specificity; examples of such models are NetTCR, pMTnet and DeepTCR [15–17]. These models are trained using labeled datasets containing known TCRs and their corresponding antigens. As a result, the models learn the TCR features that determine the specificity for a particular antigen.

Here, we introduce TouCAN, which is a contrastive learning algorithm for TCR-clustering and specificity-prediction and is trained using high-quality paired-chain TCR and labeled pMHC data. TouCAN (**T**CR clustering by **C**ontrastive learning on **An**tigen specificity) combines both distance- and feature-based strategies by utilizing contrastive learning. Originally developed for image recognition and clustering tasks [18], contrastive learning learns the distance between two objects based on their features. Contrastive learning algorithms offer the advantage of accommodating an unlimited number of classes and clustering of highly diverse data. It has been successfully applied in biological tasks such as transcription factor binding prediction[19] and enzyme function annotation [20]. Recently, contrastive learning has also been applied to TCR binding specificity prediction, however only to CDR3β sequences [21]. In addition to contrastive learning, TouCAN leverages protein language models (pLM) such as evolutionary scale modeling (ESM) [22,23] for TCR sequence embedding. pLMs are trained on large corpuses of protein sequence datasets and provide meaningful protein representations, enabling predictions of protein structure and sequence variant to function. We compare different TCR encoding methods, demonstrating superior performance of pLM-based encoding over other methods (one-hot encoding). Next, we demonstrate that TouCAN effectively clusters TCRs with the same specificity, showcasing its ability to group diverse motifs together. Finally, we assess the generalization performance of TouCAN on unseen antigen (pMHC) datasets, demonstrating generalization to an extent. Our findings highlight that TouCAN exhibits comparable performance with other methods and can successfully cluster highly diverse TCR sequences.

## Materials and Methods

### Datasets

The paired-chain TCR (TCRαβ) dataset with labeled antigen (pMHC) specificity was collected from public databases such as VDJdb, IEDB, Platypus, and McPAS-TCR [24–27]. The data contained both

TCR-pMHC sequences and were filtered as follows:

- Human and murine TCR sequences (only containing standard IUPAC amino acids),
- Human and murine pMHC sequences with known TCR specificity,
- CDR3α/β length between 10 and 23 amino acids,
- CDR3α/β loops starting with C and ending on F/Y/W,
- Exclusion of data from a study with high TCR cross-reactivity due to screening approach (10X genomics) [28],
- Peptide antigen length ranging from 8 to 15 amino acids,
- If a peptide was presented by multiple MHCs, only the MHC allele with the most TCR data was retained
- Exclusion of TCRs with known cross-reactivity
- Exclusion of peptide antigens with less than 30 specific TCRs and including a maximum number of 400 specific TCRs per antigen
- Retention of TCRs with unique CDR3β.

This resulted in 4,843 data points. To obtain a full TCRαβ sequence, TCRs without V/J gene information were removed. We used the Stitchr tool to reconstruct the full TCRαβ amino acid sequence from available V(D)J genes and CDR3αβ [29]. TCRαβ sequences that could not be reconstructed were also excluded. This resulted in 3,873 TCRαβ sequences with labeled pMHC specificity. The final dataset was clustered by 90% similarity on both CDR3α and CDR3β similarity using CD-HIT, and split into 15% validation set, 15% test set, and 70% training set (seven folds in total). Similar TCRs, as determined by CD-HIT, were assigned to the same fold. The validation set was used for hyperparameter tuning and early stopping, while the test set was used for the final performance estimation.

### TCR sequence encoding

We compared two different ways of encoding TCRs: (i) one-hot encoding and (ii) encoding with the pre-trained pLMs based on ESM.

In one-hot encoding, each amino acid is represented as an array of 20 with all zeros except a single position with ‘1’ corresponding to this amino acid. The encoded TCR results in a two-dimensional array (TCR_amino_acid_length, 20). For one-hot encoding, all six concatenated CDRα/β sequences were utilized and were concatenated with a single ‘_’ separator, except for the last CDR3α/β loop, which was padded to the maximum length.

In the case of ESM encoding, we examined two distinct models, namely ESM1-v [22] and ESM2 [23]. The ESM1-v model facilitates protein variant function prediction, whereas ESM2 encodings demonstrate state-of-the-art performance in structural predictions. We further explored diverse TCR inputs such as complete TCRαβ sequence, separate encoding of TCRα and TCRβ chains (with constant (C) domain), and individual encoding of Vα and Vβ regions (Fig. 1a). The separately encoded chains were subsequently concatenated. Notably, while the TCR amino acid length input varies, the output ESM encodings always have the same dimension through an array of 1280 numbers. Since ESM models are trained on large amounts of protein data, we anticipated that some TCR sequence-to-function properties would be captured by ESM encodings. Examination of the raw encodings for all three TCR inputs reveals that specificity clusters can be observed even in the absence of training (Fig 1b).

**Figure 1.**
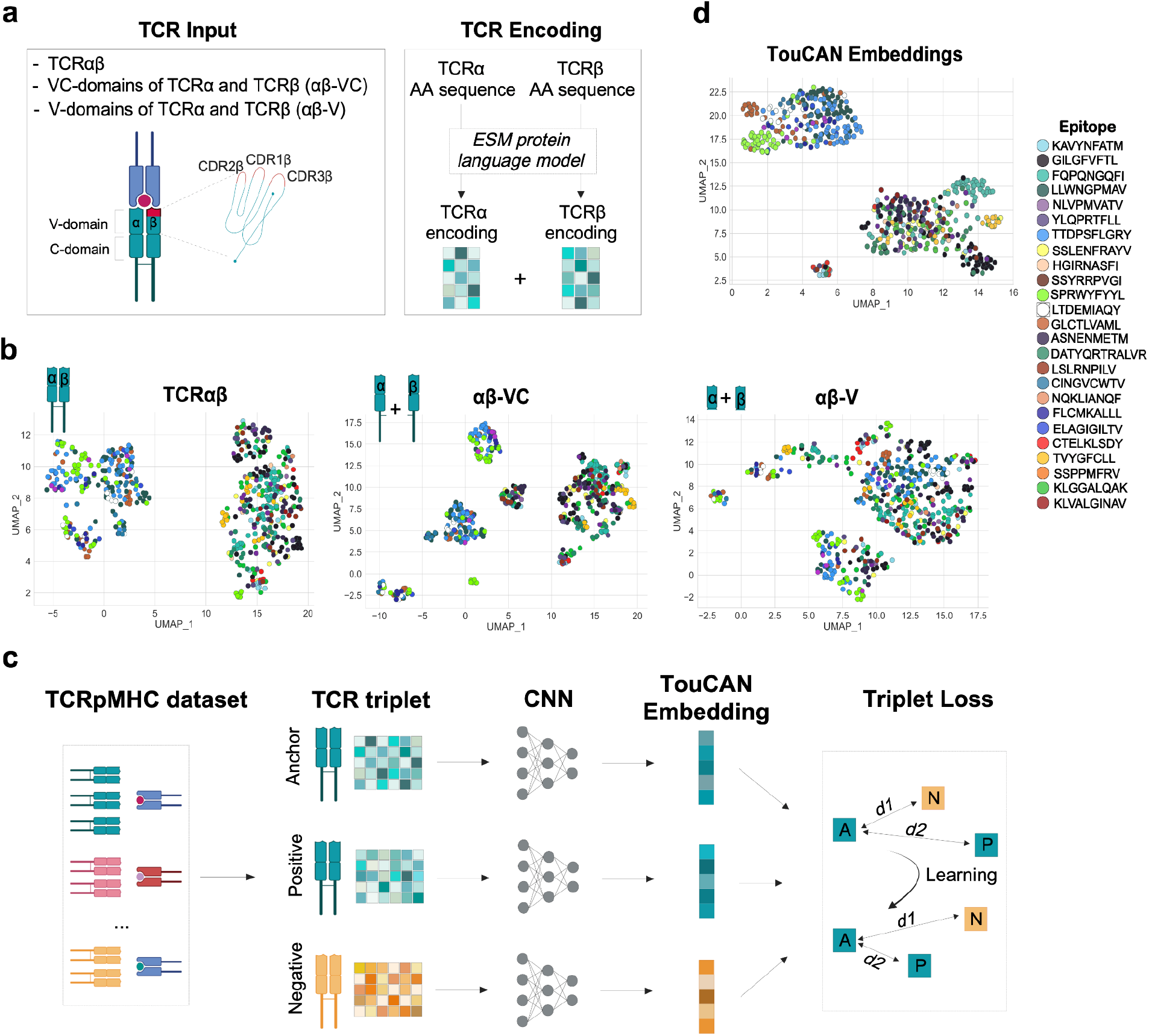
Overview of TouCAN architecture and training. **(a)** Schematic representation of a T-cell receptor (TCR) structure and TCR encoding using the ESM protein language model. **(b)** Visualization of TCR ESM encodings. The output from the ESM-1v model for various TCR input types (TCRαβ, αβ-VC, αβ-V) is visualized with UMAP, with TCR encodings color-coded by antigen (pMHC) specificity. **(c)** Schematic of TouCAN architecture shows TCR encodings are input for a CNN in triplets: ‘anchor’ and ‘positive’ TCRs target the same antigen, while a ‘negative’ TCR recognizes a different antigen. The CNN extracts a numerical embedding vector for each TCR (TouCAN embedding). Euclidean distances are calculated between the embeddings for the ‘anchor (A)’ and ‘negative (N)’ or (*d1*), and ‘anchor’ and ‘positive (P)’ or (*d2*). The network weights, and consequently the embeddings, are updated by triplet loss to minimize the distance between ‘A’ and ‘P’ and maximize the distance between ‘A’ and ‘N’ (*d1 >> d2*). **(d)** TCR embeddings learned by TouCAN visualized using UMAP and color-coded by antigen specificity. The TCR input type is αβ-V, and the encoding is obtained from ESM-1v.

To make ESM inputs more comparable with one-hot encoding and enable the use of convolutional neural networks (CNN), we reshaped the ESM output into a two-dimensional array of (64, 20) for a full TCRαβ or (128, 20) for independently encoded chains, respectively.

### TouCAN Model Architecture

TouCAN employs a contrastive triplet loss to learn TCR embeddings that represent their pMHC specificity (Fig. 1c). Initially, triplets of TCRs are selected from the training batch, with two distinct TCRs recognizing the same pMHC (serving as an anchor and a positive instance), while the third TCR recognizes a different pMHC (serving as a negative instance). A CNN is applied to these TCR triplets to extract features and acquire condensed numerical TCR embeddings. Afterwards, the Euclidean distance is calculated between the embeddings of an anchor (A) and positive (P) TCR (*d2*), as well as an anchor and negative (N) TCR instance (*d1*). To update the network weights, the triplet loss is then calculated by the following equation [18]:

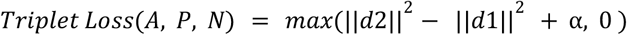

Here, α represents a margin between learned positive and negative Euclidean distances and can be adjusted accordingly.

According to the triplet loss formula, the TCR embeddings are learned in a manner that minimizes the Euclidean distance between TCRs recognizing the same antigen, while simultaneously increasing the distance for TCRs with different specificities. The learned TouCAN embeddings, colored by peptide antigen, are visualized on a test dataset (Fig. 1d).

### TouCAN training and parameter optimization

TouCAN hyperparameters were optimized using gridsearch and included learning rate, batch size, and dimensionality of the learned embeddings. All models were trained in a cross-validated manner to obtain performance distribution. The optimal model hyperparameters were selected on a validation fold and the performance of the selected model was then estimated on a test fold. Final TouCAN architecture has three convolutional layers with 128, 64m, 16 filters and ReLU activation function. The model was trained with learning rate = 0.001, batch size = 256, semi-hard triplet loss and with early stopping on a validation fold. The dimensionality of TouCAN’s output was individually optimized for each ESM model and TCR input type (Supp. Fig. 1-3).

### Clustering quality metrics

Learned TouCAN TCR embeddings were grouped into clusters by the DBSCAN algorithm using Scikit-learn Python library [30,31]. TCR cluster assignments were used to calculate three metrics: cluster retention, purity and consistency, as previously described [8]. Model comparisons were conducted based on the Purity-Retention and Purity-Consistency curves.

### CDR3 amino acid motifs

Multiple sequence alignment of correctly clustered CDR3α and CDR3β sequences was performed by MUSCLE [32]. Then, the alignment was visualized as a sequence logo by the WebLogo tool [33]

### Assigning pMHC antigen labels to TCR embeddings

The peptide antigen labels were assigned to learned TouCAN TCR embeddings using three distinct approaches: (i) K-nearest neighbors (KNN), (ii) centroid method, and (iii) maximum separation.

### KNN Method

A KNN algorithm was implemented with the Scikit-learn library [31], and was employed for antigen (pMHC) label assignment. The KNN model used a training set of TCR embeddings with known antigen labels, which were subsequently applied to the test set embeddings, with the parameter *N* neighbors equal to 5.

### Centroid Method

A ‘centroid’ embedding for each pMHC antigen was computed on the training set based on the median of all TCR embeddings corresponding to the same antigen specificity. This resulted in 25 centroids for the 25 antigens in the dataset. To assign an antigen label to a test TCR embedding, distances to all centroids were computed, and the antigen label with the smallest Euclidean distance was assigned.

### Maximum Separation Method (MaxSep)

The MaxSep method closely follows the procedure of the Centroid method, with a crucial distinction in the final step. Instead of selecting a single pMHC antigen label with the smallest Euclidean distance, MaxSep has the capacity to choose a set of antigen labels if their centroid Euclidean distances are close to one another relative to background distances to all other antigen centroids. A detailed description and implementation of the MaxSep method was previously described [20]. Here, the MaxSep method can assign up to three antigen labels to a single TCR. Any correct assignment among these antigen labels is considered a correct classification. The proportion of TCRs with two or three assigned antigen labels is available in Supplementary Table 3.

### Benchmarking on independent datasets

We obtained the dataset and model performances, including MicroAUCs, from the IMMREP benchmark study [14]. The IMMREP benchmark dataset contains negative TCRs with a negative-to-positive ratio of 5:1, generated by mismatching the TCR-pMHC pairs and random sampling from the TCR repertoire of healthy individuals [14]. These types of negative TCRs are typically not included in contrastive learning, where instances from other classes serve as negatives. To address this, an additional training step was implemented.

Initially, TouCAN was trained on positive TCR-pMHC pairs, as described previously. Then, the learned TouCAN embeddings were obtained for all TCRs in the IMMREP dataset, and their corresponding pMHC antigens were encoded using one-hot encoding. The resulting TCR and pMHC arrays were concatenated and utilized as input into a binary classifier, which was trained using the Scikit-learn library’s Multi-layer Perceptron classifier (MLPClassifier) with default parameters. The classifier’s performance was evaluated in terms of MicroAUCs and then compared with other models used in the IMMREP study [14].

Also from the same IMMREP study [14], we compiled an additional test dataset of 506 TCRs with labels for pMHC-specificity (across seven antigens), which were not present in the previous datasets used to train TouCAN. Subsequently, we predicted the embeddings of these test TCRs using TouCAN, performed clustering with DBSCAN, and calculated Purity-Retention and Consistency-Retention curves. Since TouCAN was not trained with these antigens, the model was unable to assign antigen labels. To estimate TouCAN performance, we compared it with Levenshtein distance clustering. Additionally, we established a ‘random’ baseline where TCR sequences and their labels were randomly mixed.

### Statistical Analysis

Statistical analysis was performed using the Scipy library in Python [34]. Statistical significance of the different antigen-labeling methods was determined using paired t-tests with the alternative hypothesis ‘greater’. The same statistical procedure was applied to compare different algorithms for TCR clustering.

## Results

### TCR encoding with pre-trained protein language models leads to superior performance

To assess the impact of pre-trained pLMs, we assessed the performance of TouCAN TCR clustering using ESM-based encoding vs. one-hot encoding (of all CDR loops). Additionally, simple Levenshtein Distance (LD) clustering has shown promising performance compared to more complex machine learning algorithms [8,35], therefore we use LD as an additional baseline for estimating model effectiveness. The TouCAN model underwent training using various inputs, including ESM-1v and ESM-2 encodings, and three types of TCR sequence inputs: a complete TCR sequence (TCRαβ), full alpha and beta chains encoded separately (αβ-VC), and separately encoded V-domains of the alpha and beta chains (αβ-V). This training was conducted using a cross-validation approach. Additionally, the model’s performance was evaluated and compared using Consistency-Retention and Purity-Retention metrics (Fig. 2). While the purity of TCR clusters remained unaffected by the different encoding methods, we observed a drastic difference in model consistency. All models using ESM-encoding of TCRs demonstrated improved consistency compared to one-hot encoding. Notably, separately encoded V-domains of alpha and beta chains by the ESM-1v model (αβ-V_ESM-1v) exhibited the best performance and were thus selected for subsequent analysis.

**Figure 2.**
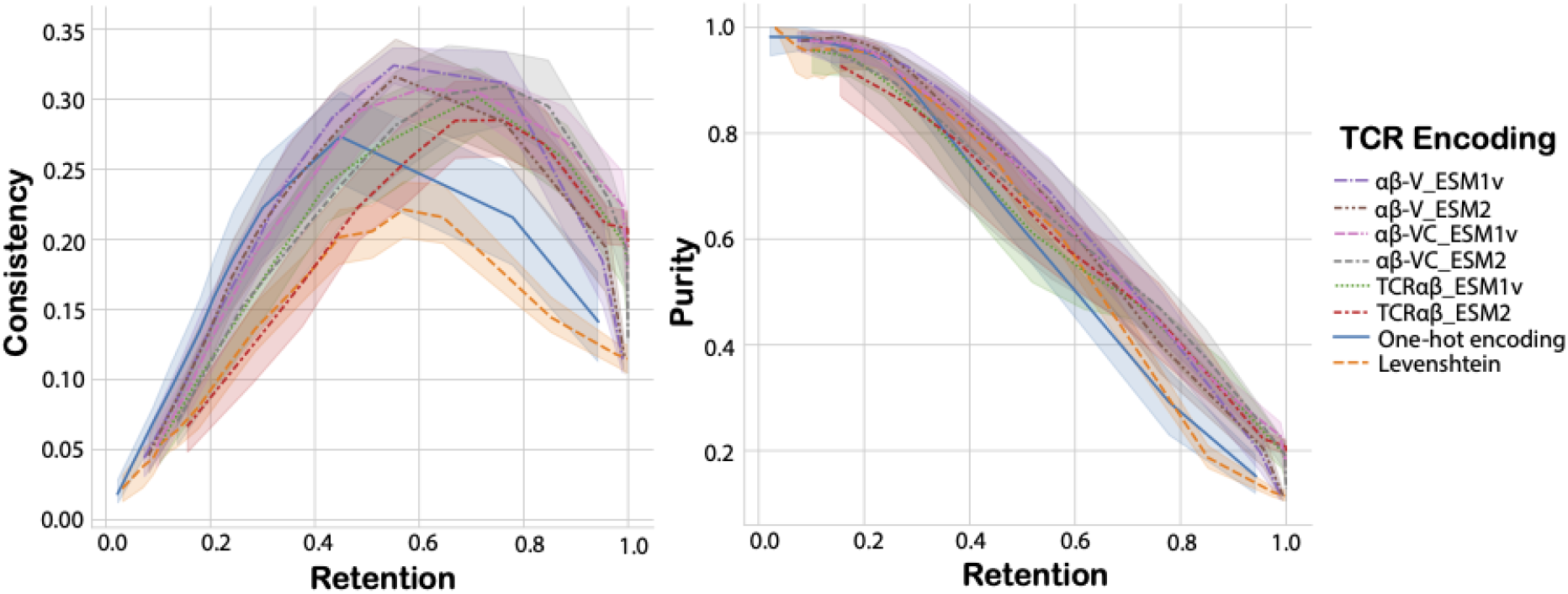
Clustering performance of different TCR inputs and encodings. Consistency-Retention and Purity-Retention curves were used to compare the clustering performance of various TCR inputs and encodings. The shaded areas indicate variation in cross-validation performance. The comparison includes TCRαβ, αβ-VC, αβ-V TCR inputs encoded by either ESM-1v or ESM2 models. One-hot encoding of CDR123αβ and Levenshtein distance serves as the baseline for comparison.

### TouCAN is able to group diverse TCR sequences into antigen-labeled clusters

Next, we analyzed TCR clusters assigned by the selected input and encoding model version of TouCAN (αβ-V_ESM-1v). TCR specificity clusters identified by TouCAN were visualized using network plots and compared with Levenshtein distance clustering at 50% dataset retention, meaning half of the test dataset was assigned to a cluster (Fig. 3a). As expected, the majority of TCR specificity clusters identified by TouCAN aligned with the most abundant antigens in the dataset (Supp. Table 2).

**Figure 3.**
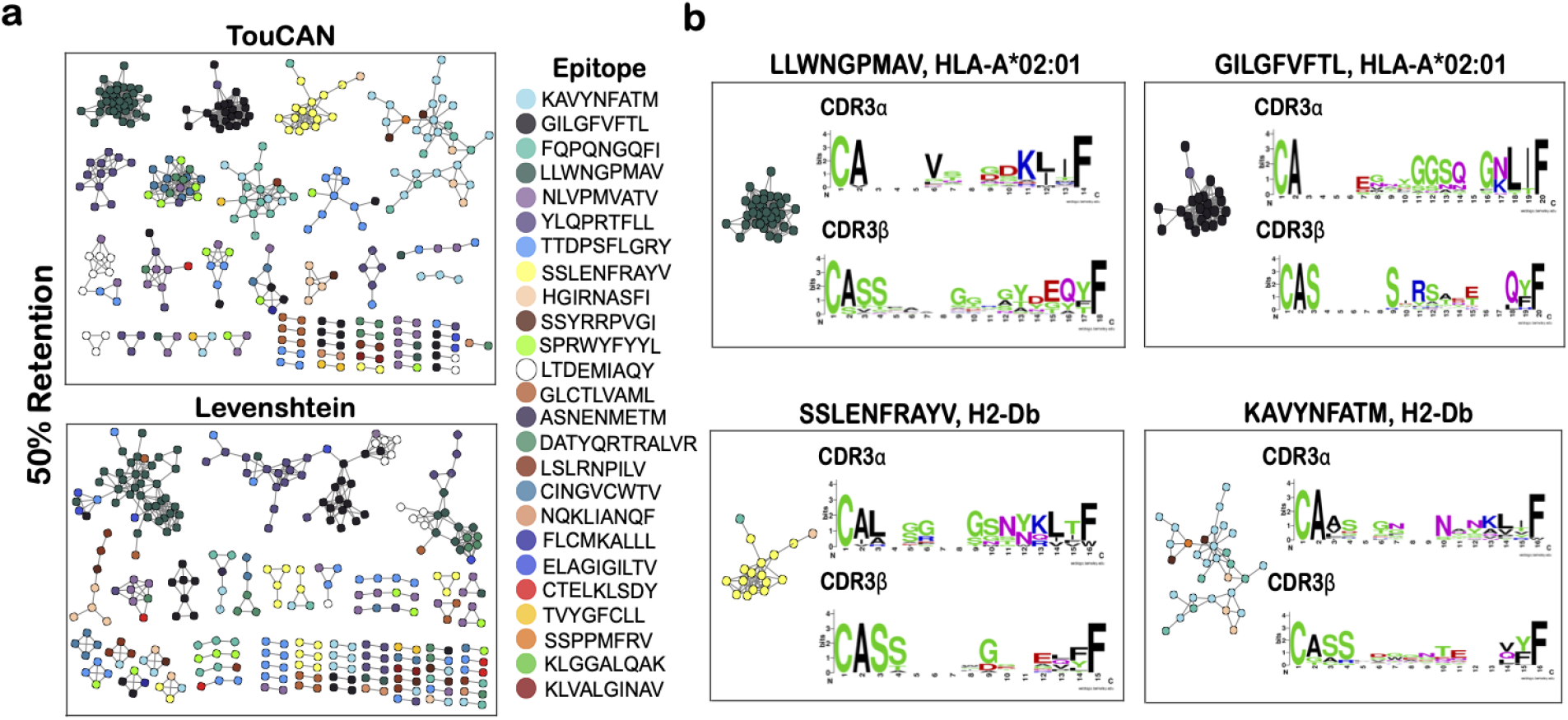
Diversity of TCR clusters identified by TouCAN. **(a)** Network plots illustrating TCRs clustered by TouCAN or Levenshtein distance at 50% test set retention. Each dot represents a TCR sequence, with TCRs color-coded by antigen (pMHC) specificity. Legend corresponds to peptide antigen sequences. **(b)** Protein sequence logos represent the largest TCR clusters identified by TouCAN. Multiple sequence alignment was performed CDR3α and CDR3β sequences of correctly assigned TCRs within a specificity cluster. The *LLWNGPMAV-HLA-A02:01* cluster contains 35 TCRs (35 correctly assigned), *GILGFVFTL-HLA-A02:01* has 23 TCRs (22 correctly assigned), *SSLENFRAYV-H2-Db* cluster consists of 17 TCRs (15 correctly assigned), and *KAVYNFATM-H2-Db* has 31 TCRs (19 correctly assigned TCRs).

TouCAN identified several TCR specificity clusters that were not captured by Levenshtein distance, highlighting the ability to assign antigen clusters to highly diverse TCR sequences. To further characterize TouCAN clusters, we performed a multiple sequence alignment of the CDR3α and CDR3β for TCRs specific to the same pMHC antigen. The resulting motifs for the four largest TouCAN clusters revealed TCRs with differing CDR3α and CDR3β lengths, as well as diverse CDR3β amino acid compositions (Fig. 3b and Supp. Table 1).

It is worth noting that ‘impure’ TCR clusters that possess several pMHC antigens consisted of either exclusively human or murine TCRs, indicating that TouCAN successfully learned to separate these two species (Supp. Table 1).

### TouCAN assignment of antigen labels to TCRs

Next, we sought to investigate the feasibility of assigning pMHC antigen labels to TCRs based on the learned TouCAN embeddings, including a comparison with different labeling approaches (Fig. 4). We implemented three labeling methods:(i) k-nearest neighbors classifier (KNN), (ii) Centroid, and (iii) Maximum Separation (MaxSep). KNN is a standard method to classify the embeddings obtained by contrastive learning [18]. Centroid and Maximum Separation methods have been successfully applied to annotate enzyme functions after contrastive learning on large-scale protein sequence data [20].

**Figure 4.**
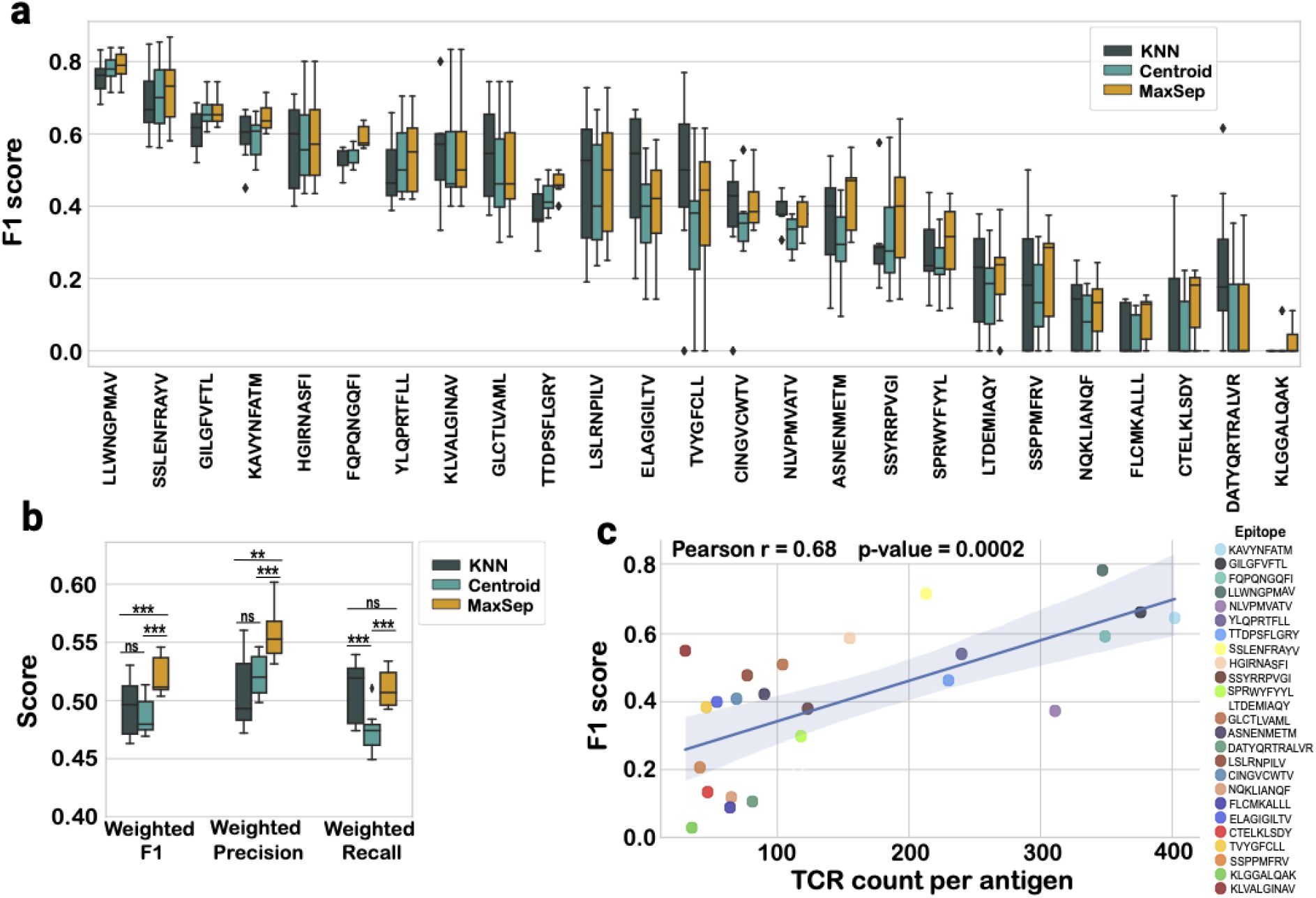
Assigning antigen labels to TCRs by TouCAN. **(a)** F1-score per antigen (pMHC) for the three antigen label assignment methods: KNN, Centroid and MaxSep. **(b)** Weighted average F1-score, Precision and Recall for antigen assignment methods calculated per fold. The weighted average takes into account the number of instances for each class, such as the number of TCRs per antigen. **(c)** Correlation between TouCAN MaxSep F1-score and the number of TCR sequences per antigen (pMHC) in the dataset. Legend indicates antigen peptide sequence per color.

The F1-score per antigen for all three labeling methods was compared in a cross-validation manner (Fig. 4a), revealing similar performance trends across all antigens. Statistical analysis revealed that the MaxSep method performed significantly better than KNN and Centroid methods (Supp. Table 4). MaxSep also performed significantly better in weighted precision and F1-score (Fig. 4b), leading us to use this approach for subsequent analysis.

Next, we investigated how TouCAN labeling performance per antigen is influenced by the abundance of TCRs specific to an antigen in the dataset (Fig. 4c), as well as the diversity of TCRs, which was measured by the average Levenshtein distance of TCRs belonging to the same antigen cluster (Supp. Fig. 5). This analysis revealed a positive correlation between F1-score and TCR count per antigen, suggesting that increased TCR data could enhance algorithm performance. Additionally, we observed a negative correlation between TouCAN performance and TCR cluster diversity, particularly on CDR3αβ Levenshtein distance (Supp. Fig. 5b), aligning with existing literature findings that higher-complexity TCR-antigen pairings are additionally challenging for predictions [14].

### TouCAN performance on unseen epitopes

TouCAN can effectively identify TCR specificity groups and assign labels for pMHC antigens present in the training dataset. However, the potential antigenic diversity space is vastly larger than what exists in current databases, therefore it is important to determine whether TouCAN can generalize to unseen antigens and group TCRs into corresponding specificity clusters. To investigate this, we curated data for 506 TCR sequences with known specificity to seven pMHC antigens. The TCR sequences were then processed through TouCAN to perform embedding and clustering.

We compared the Retention-Purity and Retention-Consistency curves of TouCAN with Levenshtein distance clustering, as well as their respective random baselines (Fig. 5a). Expectedly, both metrics outperformed the random baseline. Although Levenshtein distance clustering exhibited superior performance in cluster purity, TouCAN demonstrated higher consistency (Fig. 5a). The cluster networks for TouCAN and Levenshtein at 50% dataset retention for the unseen antigens are illustrated in Supplementary Figure 8.

**Figure 5.**
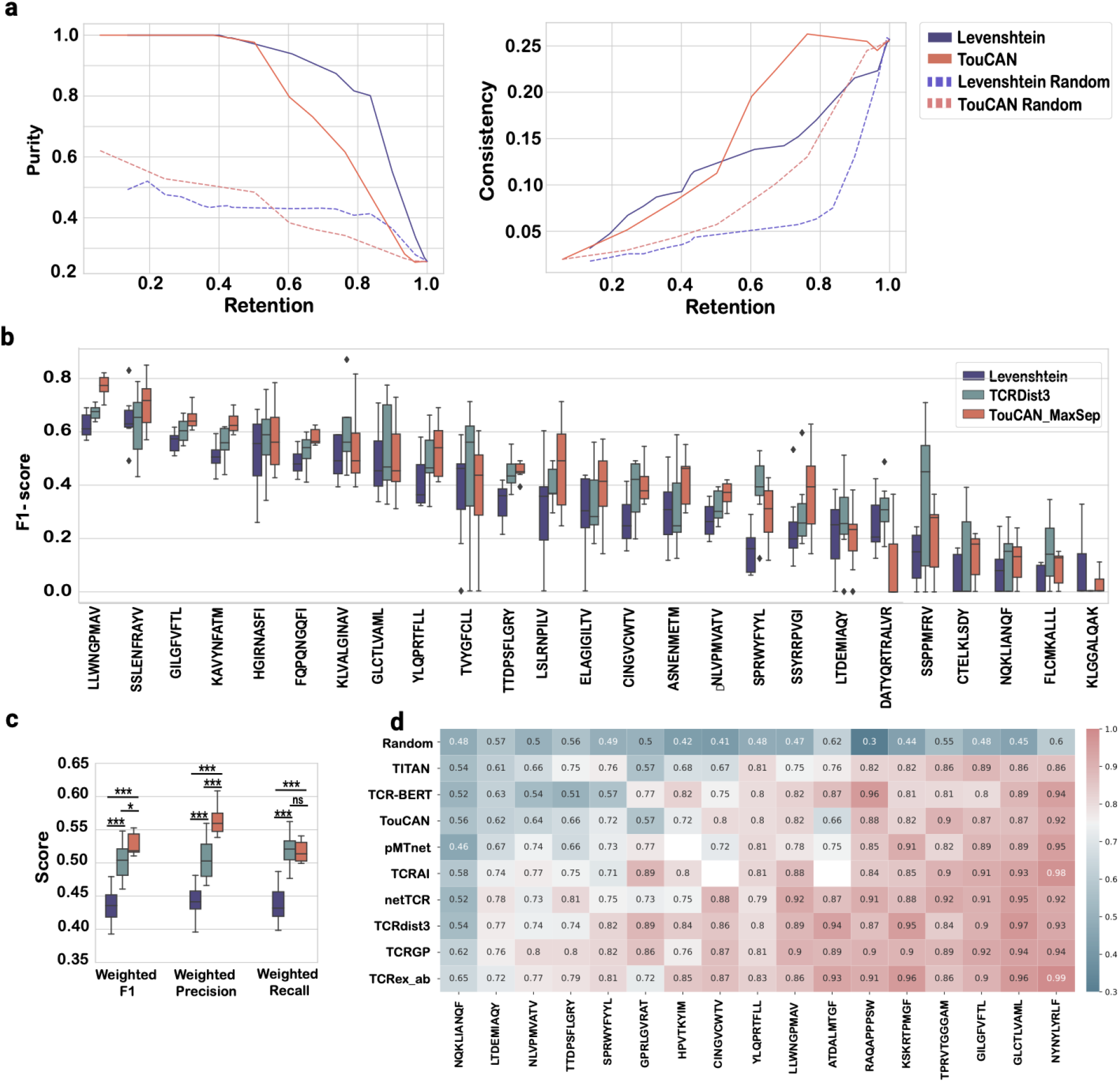
TouCAN benchmarking on unseen epitopes and state-of-the-art tools. **a**. Comparison of Purity-Retention and Consistency-Retention curves for TouCAN and Levenstein distance clustering on unseen antigens (7 unseen pMHC, 506 TCRs). Dotted lines represent a random baseline for both methods. **b**. Comparison of F1-scores per antigen for TouCAN, TCRdist3 and Levenshtein distance on the TouCAN dataset. Variation in scores indicates differences in cross-validation results. **c**. Weighted average F1-score, precision, and recall for TouCAN, TCRdist3, and Levenshtein distance on the TouCAN dataset. **d**. Heatmap of micro AUCs per antigen per method on the independent IMMREP dataset [14]. Heatmap is sorted by an increase in the average micro AUC from top to bottom.

### Comparative analysis of TouCAN with other methods

Our next objective was to compare TouCAN with existing state-of-the-art methods in TCR specificity clustering or binding prediction. While there are over 20 different algorithms available for these purposes [6,7,16,21,36–38], most of them primarily focus on the TCR-beta chain alone. Among the tools utilizing paired-chain TCR data, TCRdist3 has demonstrated state-of-the-art performance in two recent benchmarking studies. It maintained its performance even in the presence of many synthetic TCRs per antigen [35]. Additionally, the performance of TCRdist3 was not affected by the complexity of the training data [14]. We tested TCRdist3 and LD on the TouCAN dataset (Fig. 5b). Our train-test split by 90% CDR3αβ sequence similarity provided a more challenging clustering problem than frequently utilized random split. TCRdist3 was trained and tested in a cross-validation procedure identical to TouCAN. For the LD benchmark, pMHC antigen labels were assigned to TCRs in a test set based on the closest TCRαβ sequence in a train set. Both TCRdist3 and TouCAN outperformed the LD benchmark in terms of F1-score (Fig. 5b). Additionally, TouCAN showed the highest F1-score on several antigens (Fig. 5b). Further statistical analysis indicated that TouCAN and TCRdist3 performed comparably to one another (Suppl. Table 5). When we compared weighted F1-score, Precision and Recall scores, TouCAN exhibited a higher precision than TCRdist3 and LD, however, both TouCAN and TCRdist3 methods had similar recall scores (Fig. 5c).

Finally, we evaluated TouCAN’s performance on an independent benchmarking dataset from the IMMREP study [14]. To achieve this, TouCAN was re-trained on the IMMREP dataset (as described in the Materials and Methods ) and micro AUCs were compared per method and per antigen. In this analysis, TouCAN demonstrated performance on par with other tools but did not exhibit superior performance over all of the available methods (Fig. 5d). This observation may be attributed to the random split of the IMMREP dataset, where TCR sequence similarity is higher between the train and test sets, making it easier to identify TCRs binding the same antigen. Thus, TouCAN may provide an advantage in scenarios where TCR sequences sharing specificity to the same antigen possess sequences that are more dissimilar to each other.

## Discussion

Identifying the antigen (pMHC) specificity of TCRs is an essential criteria for the development of safe and targeted T-cell therapies [39], as well as facilitating progress in other TCR-based therapeutics (e.g., soluble TCR biologics) [40]. While experimental methods for discovering TCR specificity are advancing, they still require extensive molecular and cellular engineering as well as screening efforts [11], underscoring the need for computational approaches to predict TCR-antigen specificity [10]. TCR clustering for antigen-specificity predictions offers one powerful approach. Various methods have been developed for sequence-based TCR clustering that are able to produce high purity. However, TCRs that share a common antigen specificity can have highly dissimilar Vα and Vβ sequences. Therefore, such clustering methods often struggle to group such diverse TCRs into a single antigen-specific cluster. Contrastive learning algorithms excel in clustering highly diverse datasets by learning distances between input objects, such an approach has recently been applied on large scale protein sequence data for enzyme function annotation [20]. Concurrently, the development of large language models for proteins has led to important advancements in protein structure, function and evolution predictions [23,41,42]. While several studies have applied TCR-specific pLMs to TCR prediction [38,43], extensive research on the application of general pLMs, such as ESM, is lacking.

In this study, we explore the viability of utilizing contrastive learning and ESM embeddings for TCR clustering and antigen specificity applications. Our findings demonstrate a significant enhancement in clustering performance when employing TCR encoding by ESM, particularly in terms of consistency. Encoding TCR V-domains with ESM-1v yielded the highest clustering performance based on consistency and retention metrics. Additionally, we observed that while raw ESM embeddings alone exhibit a certain degree of TCR clustering, the incorporation of contrastive learning refines and expands these clusters.

Furthermore, we highlight the capability of TouCAN in grouping TCRs with diverse CDR3α and CDR3β sequences, as well as varying CDR3 lengths, resulting in improved cluster consistency. Notably, TouCAN did not cluster murine and human TCR sequences together, indicating its ability to distinguish TCR species based on V-domain sequence information. Finally, we demonstrate that assignment of TCR-antigen labels based on TouCAN embeddings results in comparable performance with state-of-the-art methods in the field such as TCRdist3.

### Limitations to the Study

TouCAN was trained on paired-chain high-quality TCR data, which, while providing better performance, such data remains scarcely available. Our study highlighted the dependence of TouCAN’s performance on antigen abundance in the dataset. We anticipate improvement in performance as the dataset size increases. Additionally, TCR cross-reactivity was not addressed in this study, as cross-reactive TCRs were excluded from the dataset. With growing TCR-pMHC datasets, incorporating cross-reactivity into training becomes more important. The implemented MaxSep method can accommodate labeling of cross-reactive TCRs by allowing for the assignment of multiple antigens, although its performance on validated cross-reactive TCRs was not tested. Lastly, while TouCAN exhibits the ability to generalize to unseen antigens, its performance is comparable to simple Levenshtein distance clustering. Addressing these limitations will be crucial for the continued development and effectiveness of TCR-clustering and specificity prediction methods.

## Supporting information

TouCAN supplementary

## Authors’ Contributions Statement

M.P and S.T.R conceived the main idea and the framework of the manuscript. M.P, O.F and D.S. collected the data, M.P performed the machine learning analysis. M.P and S.T.R wrote the manuscript, with input from all other co-authors.

## Data and Code Availability

The data and code used to perform the work in this study will be available at the following: https://github.com/LSSI-ETH/TouCAN

