## Supplementary material for "TCR clustering by contrastive learning on antigen specificity": TouCAN supplementary: TouCAN_supplementary.pdf

### Supplementary Materials

#### Supplemental Figures

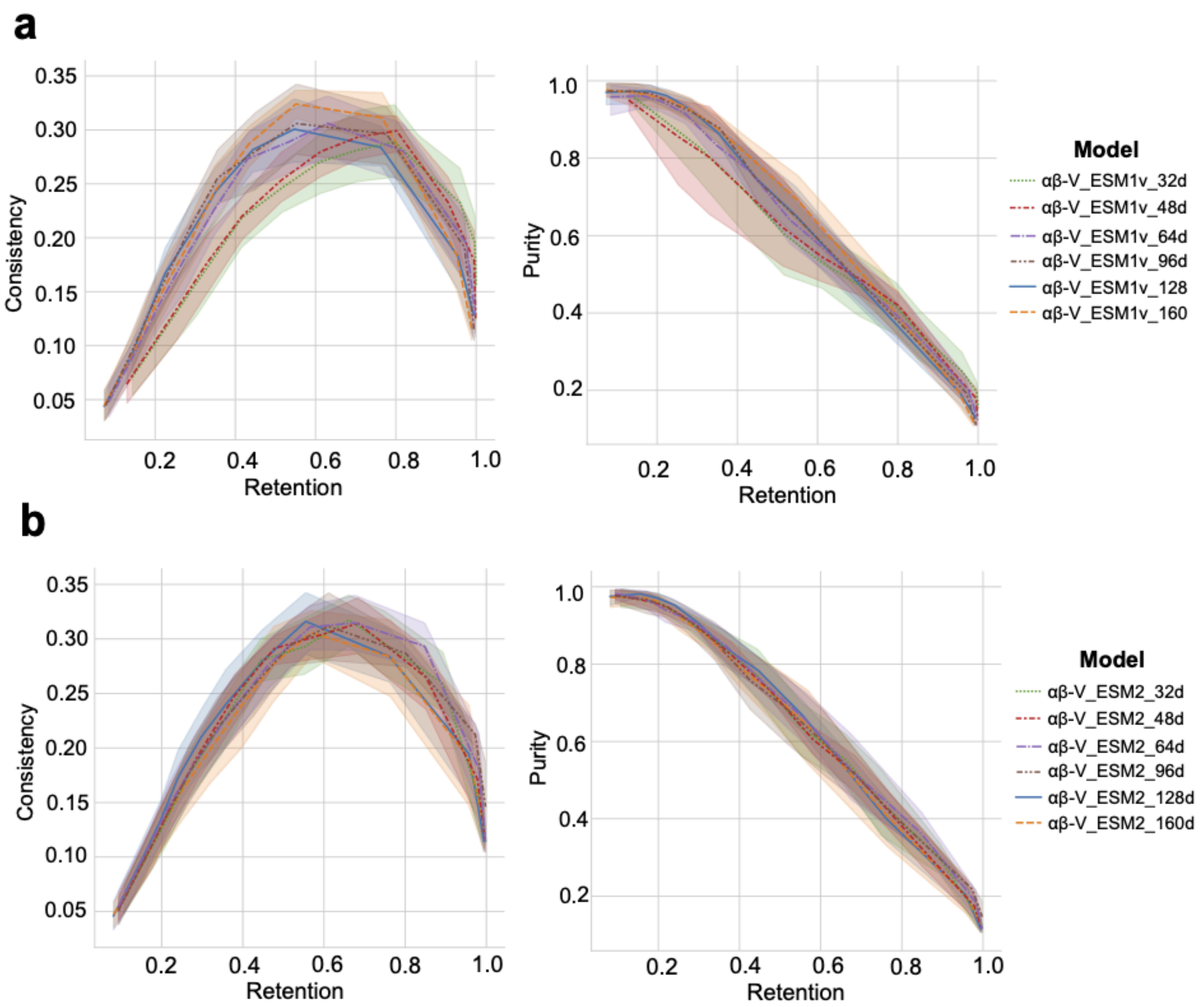


**Figure S1. Selection of the optimal TouCAN embedding dimensionality for the TCR input type 'αβ-V'. a.** Consistency-Retention and Purity-Retention curves for the models utilizing the 'αβ-V' input type encoded by ESM-1v. **b.** Consistency-Retention and Purity-Retention curves for the models utilizing the 'αβ-V' input type encoded by ESM-2. The shaded areas indicate variation in cross-validation performance.

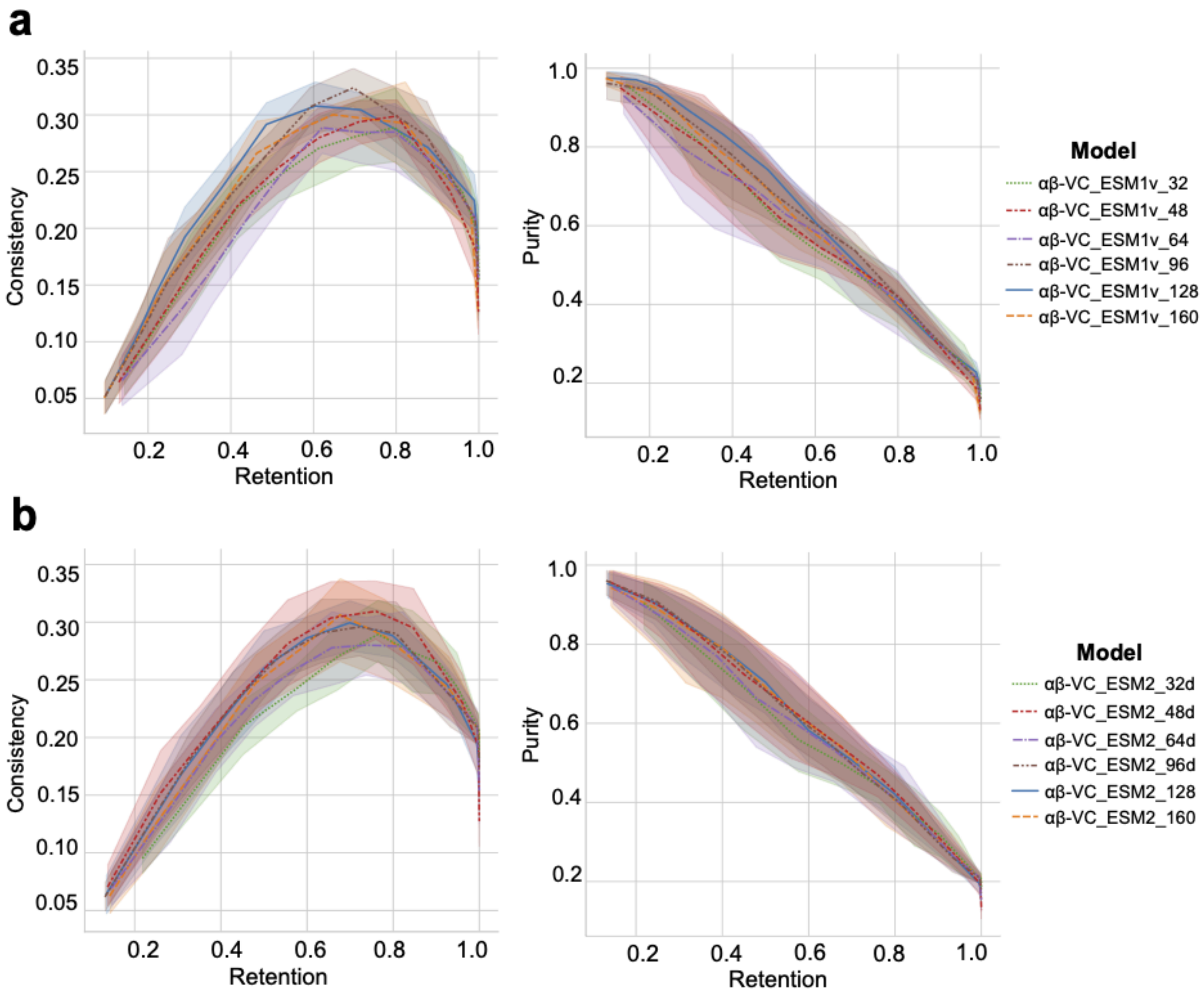


**Figure S2. Selection of the optimal TouCAN embedding dimensionality for the TCR input type 'αβ-VC'**. **a.** Consistency-Retention and Purity-Retention curves for the models utilizing the 'αβ-VC' input type encoded by ESM-1v. **b.** Consistency-Retention and Purity-Retention curves for the models utilizing the 'αβ-VC' input type encoded by ESM-2. The shaded areas indicate variation in cross-validation performance.

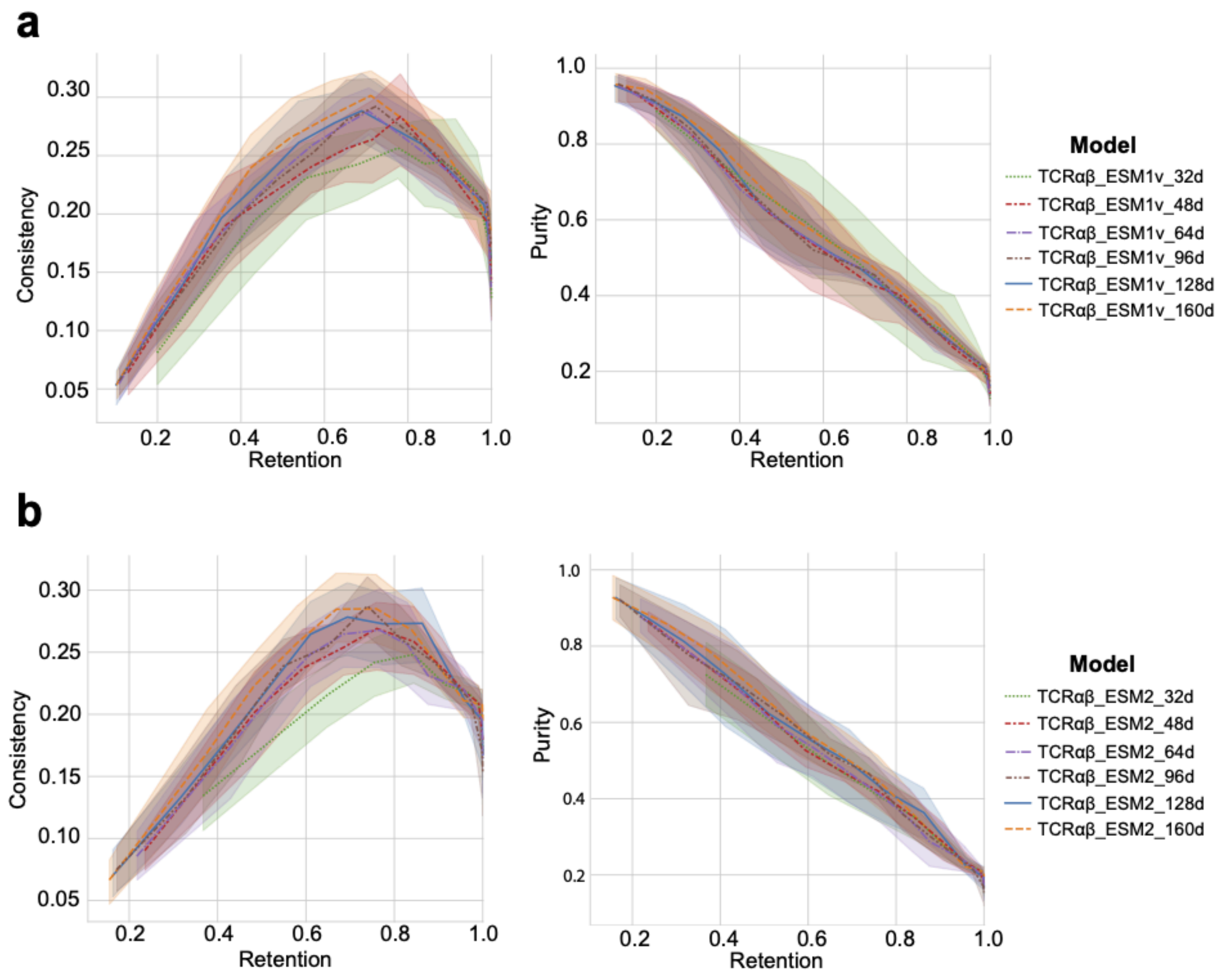


**Figure S3. Selection of the optimal TouCAN embedding dimensionality for the TCR input type 'TCRαβ'**. **a.** Consistency-Retention and Purity-Retention curves for the models utilizing the 'TCRαβ' input type encoded by ESM-1v. **b.** Consistency-Retention and Purity-Retention curves for the models utilizing the 'TCRαβ' input type encoded by ESM-2. The shaded areas indicate variation in cross-validation performance.

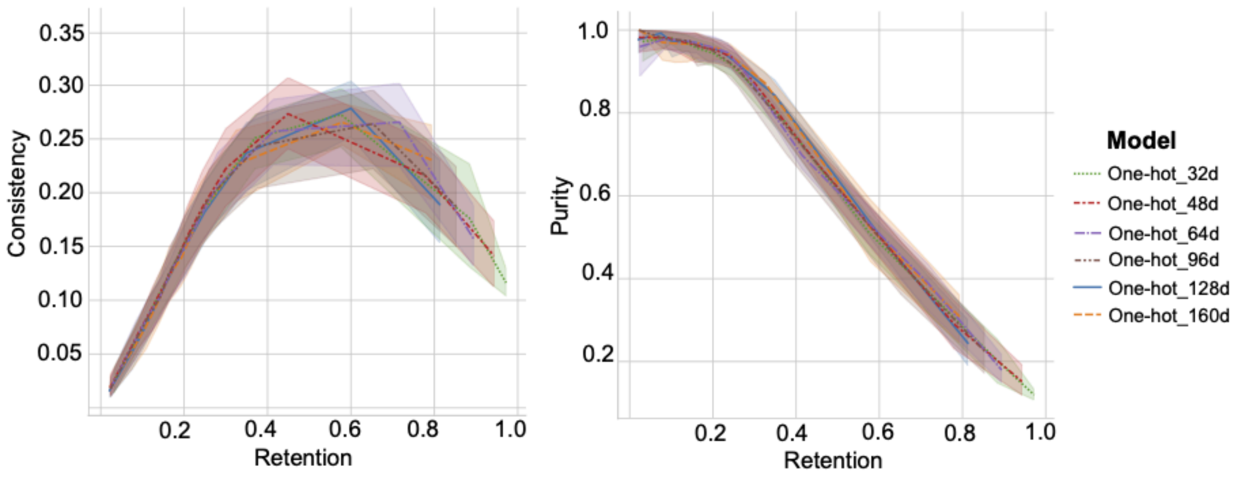


**Figure S4. Selection of the optimal TouCAN embedding dimensionality for the TCR input type ‘One-hot’**. Consistency-Retention and Purity-Retention curves for the models with concatenated CDR1-2-3αβ sequences encoded by one-hot method. The shaded areas indicate variation in cross-validation performance.

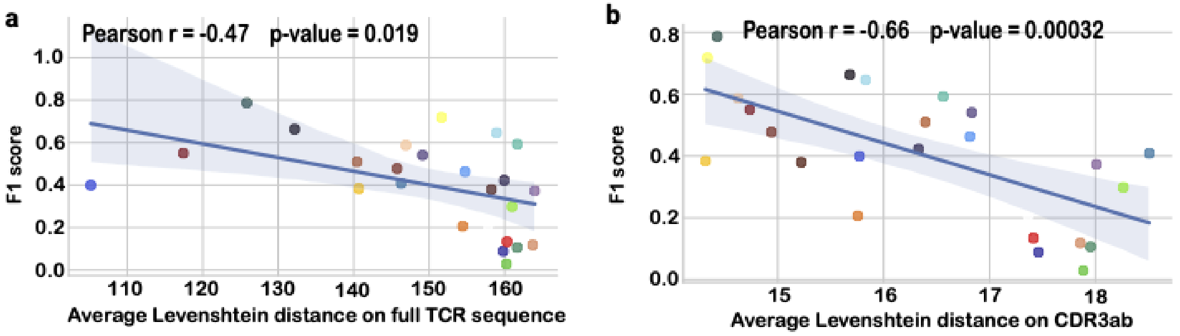


**Figure S5.** Correlation between TouCAN F1-score performance and TCR diversity per antigen calculated as average Levenshtein distances between TCRs specific to the same antigen.

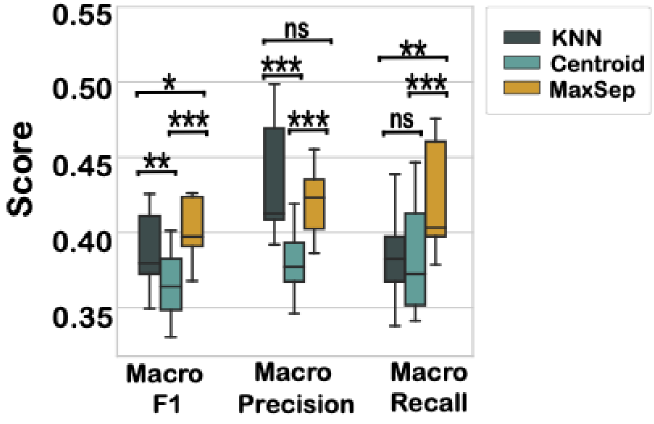


**Figure S6**. Macro average of F1-score, Precision and Recall (summed for all antigens) for different antigen label assignment methods on TouCAN dataset.

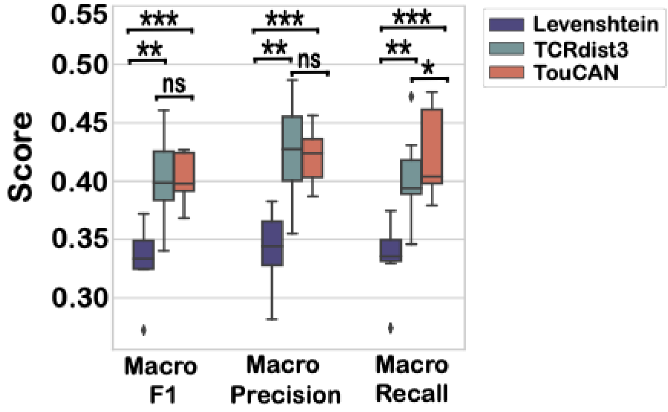


**Figure S7**. Macro average of F1-score, Precision and Recall (summed for all antigens) for different algorithms on TouCAN dataset.

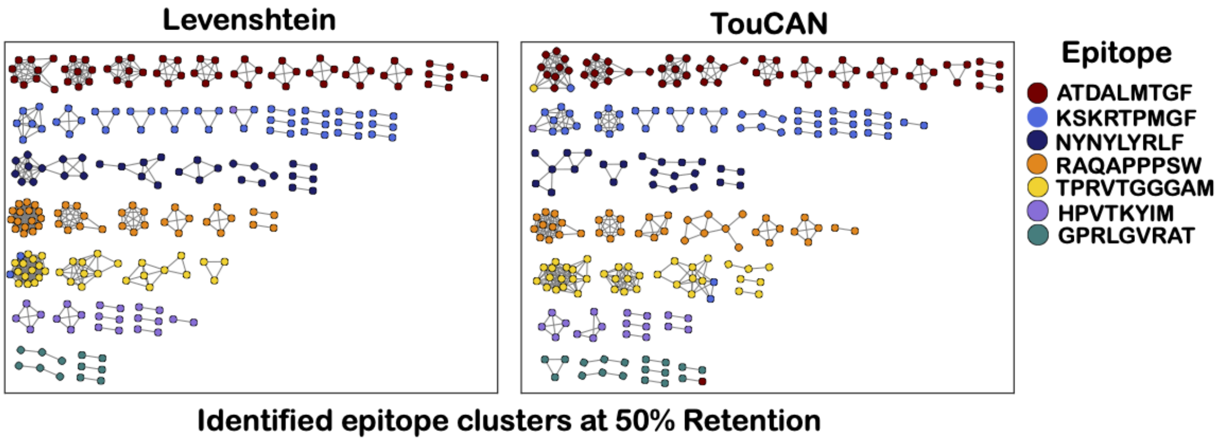


**Figure S8.** Identified TCR clusters by Levenshtein distance and TouCAN on ‘unseen’ antigens from IMMREP dataset.

### Supplemental Tables

**Supplementary Table 1.** TCR sequence information in clusters identified by TouCAN. Due to its size, this table is included as a separate csv file

**Supplementary Table 2.** TCR count per antigen in the TouCAN dataset.

| Epitope | TCRs in a dataset |
|---|---|
| KAVYNFATM | 402 |
| GILGFVFTL | 376 |
| FQPQNGQFI | 349 |
| LLWNGPMAV | 347 |
| NLVPMVATV | 311 |
| YLQPRTFLL | 240 |
| TTDPSFLGRY | 230 |
| SSLENFRAYV | 213 |
| HGIRNASFI | 155 |
| SSYRRPVGI | 123 |
| SPRWYFYYL | 118 |
| LTDEMIAQY | 116 |
| GLCTLVAML | 104 |
| ASNENMETM | 90 |
| DATYQRTRALVR | 81 |
| LSLRNPILV | 77 |
| CINGVCWTV | 69 |
| NQKLIANQF | 65 |
| FLCMKALLL | 64 |
| ELAGIGILTV | 54 |
| CTELKLSDY | 47 |
| TVYGFCLL | 46 |
| SSPPMFRV | 41 |
| KLGGALQAK | 35 |

| Epitope | TCRs in a dataset |
|---|---|
| KLVALGINAV | 30 |

**Supplementary Table 3.** The count of TCRs, which were assigned multiple antigen labels by MaxSep method.

| TCR count with multi-epitope assignment | |
|---|---|
| Second Epitope label | 662 TCRs (17,5 % of all dataset) |
| Third Epitope label | 244 TCRs (6,4 % of all dataset) |

**Supplementary Table 4.** P-values of the paired t-test on F1-score values per epitope per fold.

| Epitope assigning methods | P-value of the paired t-test, alternative = 'greater' |
|---|---|
| MaxSep vs Centroid | p-value = 1.5649e-29 |
| MaxSep vs KNN | p-value = 0.0003 |
| KNN vs Centroid | p-value = 3.8067e-09 |

**Supplementary Table 5.** P-values of the paired t-test on F1-score values per epitope per fold.

| TCR clustering algorithm | P-value of the paired t-test, alternative = 'greater' |
|---|---|
| TouCAN vs TCRdist3 | p-value = 0.4823 |
| TouCAN vs Levenshtein | p-value = 1.3147e-38 |
| TCRdist3 vs Levenshtein | p-value = 0.0012 |
